# A smartphone app for individual anesthetic calculation decreased anesthesia-related mortality in mice

**DOI:** 10.1101/2020.09.09.289728

**Authors:** Carlos Poblete Jara, Rodrigo S Carraro, Ariane Zanesco, Beatriz Andrade, Karina Moreira, Guilherme Nogueira, Bruno L. Souza, Thais Paulino Prado, Valeria Póvoa, Licio A. Velloso, Eliana P. Araújo

## Abstract

Annually, millions of animals are used for experimental purposes. Despite the recommended anesthetic doses being well-known worldwide, the final amounts applied to mice could be different than those calculated. Here, we developed, tested, and validated a mobile app where researchers and operators were able to use personal devices to process body weight, calculate a master anesthetic cocktail, and then apply the individual volume to each mouse. Our objective was to refine anesthesia procedures using information technologies. Our data showed that the “Labinsane” mobile app decreased anesthetic-related deaths upon using weight-adjusted doses of ketamine and xylazine. Also, we validated that the Labinsane mobile app matched all calculations of anesthetic doses. To our knowledge, this is the first report with hundreds of anesthetized mice records and validation and implementation of a mobile app to solve an old but transversal challenge for researchers working with experimental mice.

## Introduction

Annually, millions of animals are used for experimental purposes ^1^. Experimental mice are usually anesthetized intraperitoneally with a ketamine and xylazine solution ^2–5^. An intraperitoneal procedure permits rapid application and fast anesthetic effect. However, previous intraperitoneal ketamine and xylazine combinations have caused several challenges related to a low margin of safety, prolonged recovery and persistence of lost reflexes in mice ^4,6,7^. Ketamine and xylazine doses range from 60 to 200 mg/kg of ketamine and 4 to 26 mg/kg of xylazine ^2,4,6–9^. However, serious inconsistent rates of anesthesia-related mortality (0%-100%) have been reported ^4,6–9^.

Recent studies have shown that the MIT App Inventor, a free and open-source software, can improve the performance and quality of the data analyzed, helping operators to make well-informed decisions ^10^. A previous report validated a smartphone app for calculation of CO_2_ in inhalational anesthesia ^11^, supporting the idea that a mobile app could help operators to calculate individual anesthetic doses. Here, we developed and validated a mobile app to help researchers improve the accuracy of recommended intraperitoneal anesthetic doses applied in experimental mice. We used a range of previously tested safe doses ^4,9^, which are now contained in the “Labinsane” mobile app.

## Methods

For this study, records from C57BL/6J and Swiss mice were included. We collected anesthetic procedures described in electronic or physical handwritten documents retrospectively between 2015 and 2020 from eight researchers. Records included from different projects were approved by the Ethical Committee of the University of Campinas (4330-1A, 5521-1, 5425-1, 5349-1, 4637-1, 4072-1, 3826-1, 5414-1, 4930-1, and 4699-1). Records of anesthetic procedures were conducted according to the “Guide for the Care and Use of Laboratory Animals” ^12^.

### Criteria for inclusion

We collected the primary anesthetic records of intraoperative and anesthesia-related mortality for 8-to-16 weeks old C57BL/6J and Swiss mice. Anesthesia-related mortality was defined as lost breath or rigor mortis occurring within 2 hours after induction of anesthesia. Anesthetic procedure characteristics were as follows: intraperitoneal route, using an insulin syringe, 31G needle, sterile saline solution or distilled water for injections, 10 g/100 ml stock ketamine and 2 g/100 ml stock xylazine (C57BL/6J) or 10 g/100 ml stock ketamine, 2 g/100 ml stock xylazine and 0.5 g/100 ml stock diazepam (Swiss). Animals were on a chow diet or high-fat diet (45% fat), and individual weights were measured on the same day of the anesthetic procedures. All incomplete records were excluded from the analysis.

### Anesthetic procedure screened

Anesthetic injections are a routine procedure in our lab. We selected records with the following pattern: using only a mixture of xylazine (Anasedan, Brazil), ketamine (Dopalen, Brazil), and saline 0.9% solution. Intraperitoneal injections were performed in mice in the dorsal recumbent position. The anesthetic combination was made up as a single injection.

For both the standard dose and Individual dose, animals were weighed on the same day of the anesthetic procedure. The anesthetic cocktail was freshly prepared for each experiment, as previously described ^8^. The final solution was used immediately. Standard doses were prepared as previously described ^5^, with some local adaptations. Briefly, the final solution of the standard dose was prepared with 400 μL xylazine (2 g/100 ml), 400 μL ketamine (10 g/100 ml) and 200 μL saline solution. The standard dose cocktail (80 to 100 μL of the mixture, according to the weight) was administered intraperitoneally once by procedure. The final solution of the Individual dose was prepared and calculated by the Labinsane mobile app. Briefly, the Labinsane mobile app processes body weight, calculates a master anesthetic cocktail, and then indicates the individual volume (in μL) to administer for each individual mouse (Figure 1). A specific individual dose adjusted to the body weight of the mouse was administered intraperitoneally once by procedure. For the C57BL/6 strain, we used the range of previously tested safe doses (3, 7), and for the Swiss strain, we added diazepam (5 mg/kg) to provide remarkable sedation and full relaxation (12).

**Figure 1.**
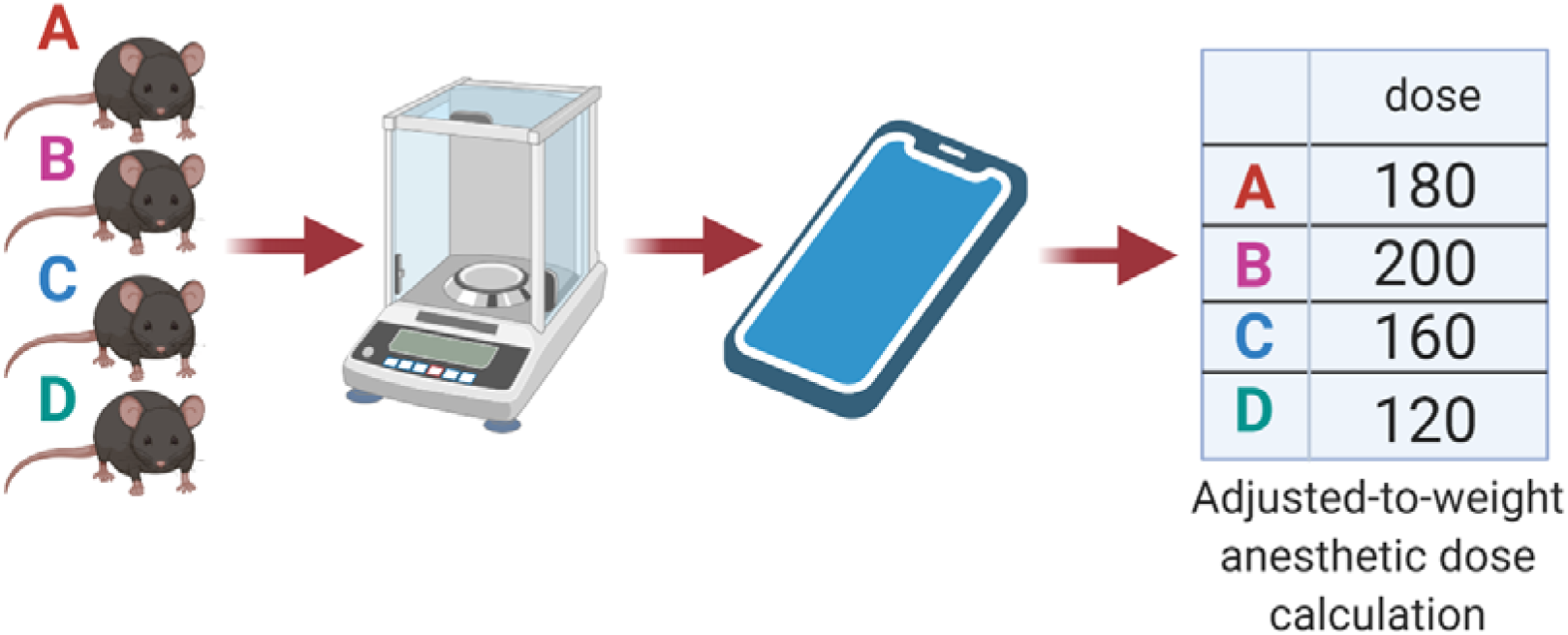
Labinsane schematic workflow. The animals are individually weighed, and measures processed by the Labinsane mobile app. A master anesthetic cocktail is calculated according to the total number of animals and individual doses are defined.

### Labinsane formula

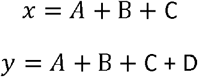

The *x* formula describes the final master anesthetic solution for the C57BL/6 mice. The *y* formula describes the final master anesthetic solution for the Swiss mice. In both formulas, A represents the final ketamine stock volume, B the final xylazine stock volume, C the final saline volume and D the final diazepam stock volume.

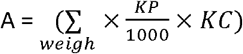

“A” from the Labinsane formula is the sum of all body weights of mice times ketamine prescription times ketamine stock concentration.

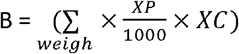

“B” from the Labinsane formula is the sum of all body weights of mice times xylazine prescription times xylazine stock concentration.

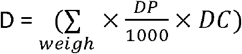

“D” from the Labinsane formula is the sum of all body weights of mice times diazepam prescription times diazepam stock concentration.

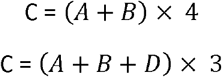

“C” from the Labinsane formula is the sum of the final ketamine stock volume and final xylazine stock volume times the dilution factor. The dilution factor here is a constant: “4” for C57BL/6 mice and “3” for Swiss mice.

The records collected described three different dose prescriptions of ketamine (mg/kg) and xylazine (mg/kg) in the following proportions: 70/7, 80/8, and 100/10, with the same prescription for diazepam (5 mg/kg). The anesthetic dose prescription was defined according to the need for long or short procedures.

### Data analysis

We pooled the data extracted from the original records on a Microsoft Excel spreadsheet (Microsoft Corporation, 2007). All statistical tests were performed in GraphPad Prism 6. Data were analyzed with Fisher’s exact test (two-sides) and odds ratios (ORs). Statistical significance was set as alpha < 0.05 and a 95% confidence interval (CI).

### Data availability

Labinsane free code will be available after publication on the Labinsane GitHub (https://github.com/blinkeado/labinsane)

## Results

### Weight-adjusted anesthetic dose decreased anesthesia-related mortality

Over the 5 years of the study, 806 anesthetized mice were evaluated, 783 on a chow diet (Table 1) and 23 mice on a high-fat diet. We obtained data from the anesthetic procedure records of the standard anesthetic dose mice and applied the weight-adjusted doses. We identified 24 intraoperative and anesthesia-related deaths within all anesthetic procedures. The Individual (weight-adjusted) dose group showed an anesthesia-related mortality rate of 1.1%. On the other hand, mice anesthetized using the standard protocol showed an anesthesia-related mortality rate of 9.3% (Table 3). The association between the weight-adjusted dose protocol and survival outcome was statistically significant (Table 3, p < 0.01). Moreover, the weight-adjusted dose procedure showed a protective effect, decreasing the ORs of both intraoperative and anesthesia-related deaths compared to the standard dose procedure (OR 0.1, 95% CI 0.04-0.24, p < 0.01).

**Table 1.**
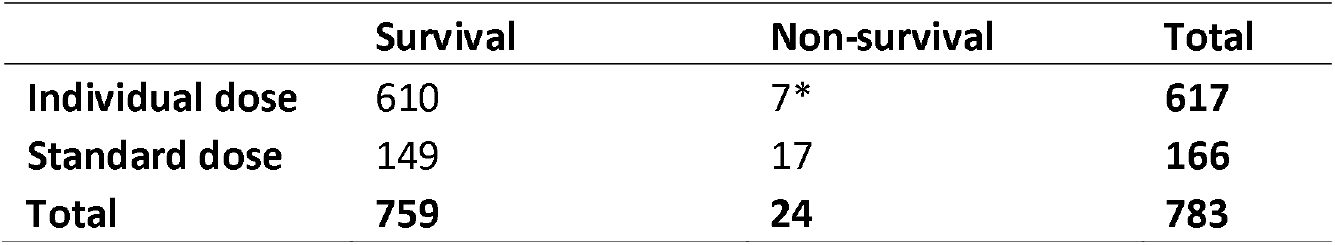
Contingency table. Survival and non-survival numbers crossed by Individual dose (weight-adjusted dose) and standard dose in Chow diet animals. Fisher’s exact test. * p < 0.05

We evaluated three weight-adjusted doses of ketamine/xylazine for short (70/7 and 80/8 mg/mg per kg) or long (100/10 mg/mg per kg) procedures. We identified higher anesthesia-related mortality with the 100 mg of ketamine/10 mg of xylazine dose compared to the other weight-adjusted doses (Table 3).

Next, we evaluated if two different diets, the chow diet (Chow) and high-fat diet (HFD), could influence the anesthesia-related mortality rate. In this way, we identified that animals fed a HFD showed the highest anesthesia-related mortality (26.1%). Indeed, we identified that animals fed a HFD were 11.1 times more likely to suffer intraoperative and anesthesia-related deaths compared to animals fed with Chow (OR 11.1, 95% CI 4.03-30.78, p < 0.01). Also, mice fed a HFD showed higher body weight compared to mice fed with Chow (Figure 2).

**Figure 2.**
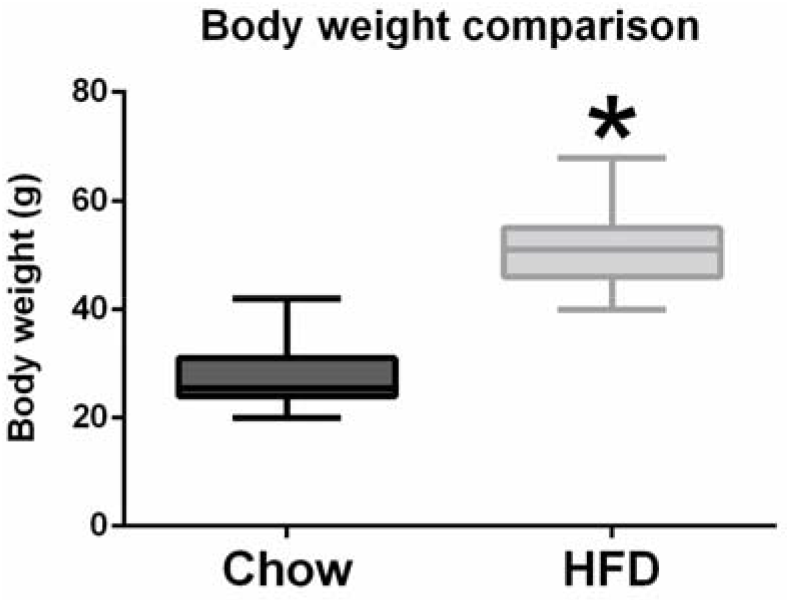
Body weight comparison. Body weight records from mice fed a chow diet (Chow) and high-fat diet (HFD). * p < 0.05

We evaluated if different strains of mice, C57BL/6 or Swiss, could influence the anesthesia-related mortality rate. We identified no significant difference in anesthesia-related mortality between C57BL/6 and Swiss mice.

### Labinsane mobile app matched all weight-adjusted anesthetic doses calculation

To validate the Labinsane weight-adjusted anesthetic doses calculation, we challenged the Labinsane mobile app to reproduce the results of individual doses calculated by a Microsoft Excel formula. We chose a Microsoft Excel formula because it is one of the top stable software available. Randomly, we tested 449 mice, and we identified that the Labinsane mobile app matched all (100%) the individual anesthetic doses calculated by the Microsoft Excel formula (Table 2) with no statistical difference (p = 0.9). These results suggested no errors in the Labinsane mobile app code.

**Table 2.**
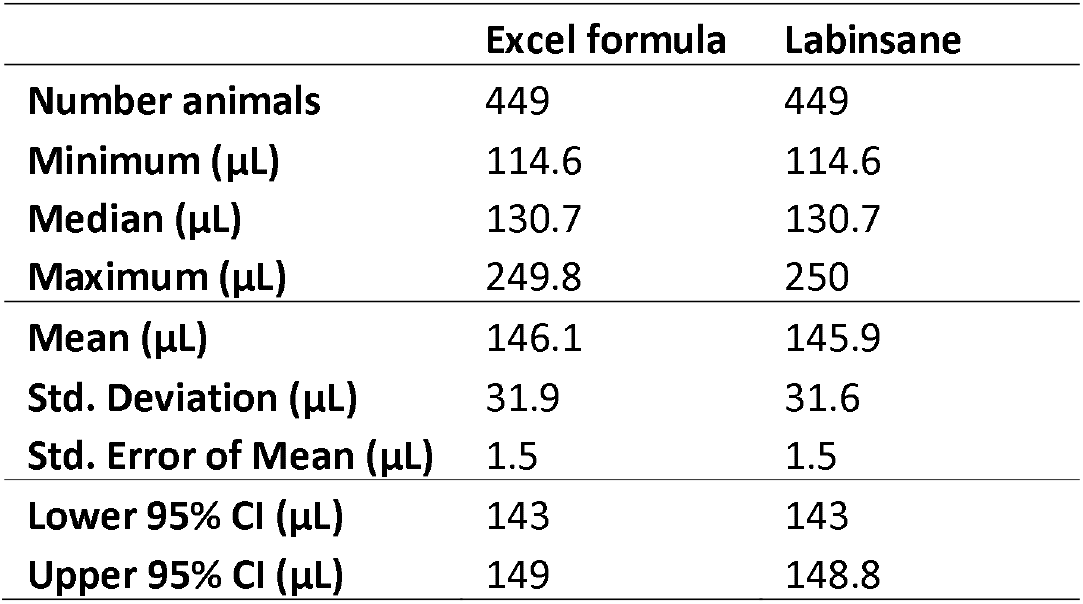
Labinsane validation. Median, mean, standard deviation, standard error and confidence interval comparing the Microsoft Excel spreadsheet and Labinsane mobile app. Unpaired t test, p value: 0.9432.

**Table 3.**
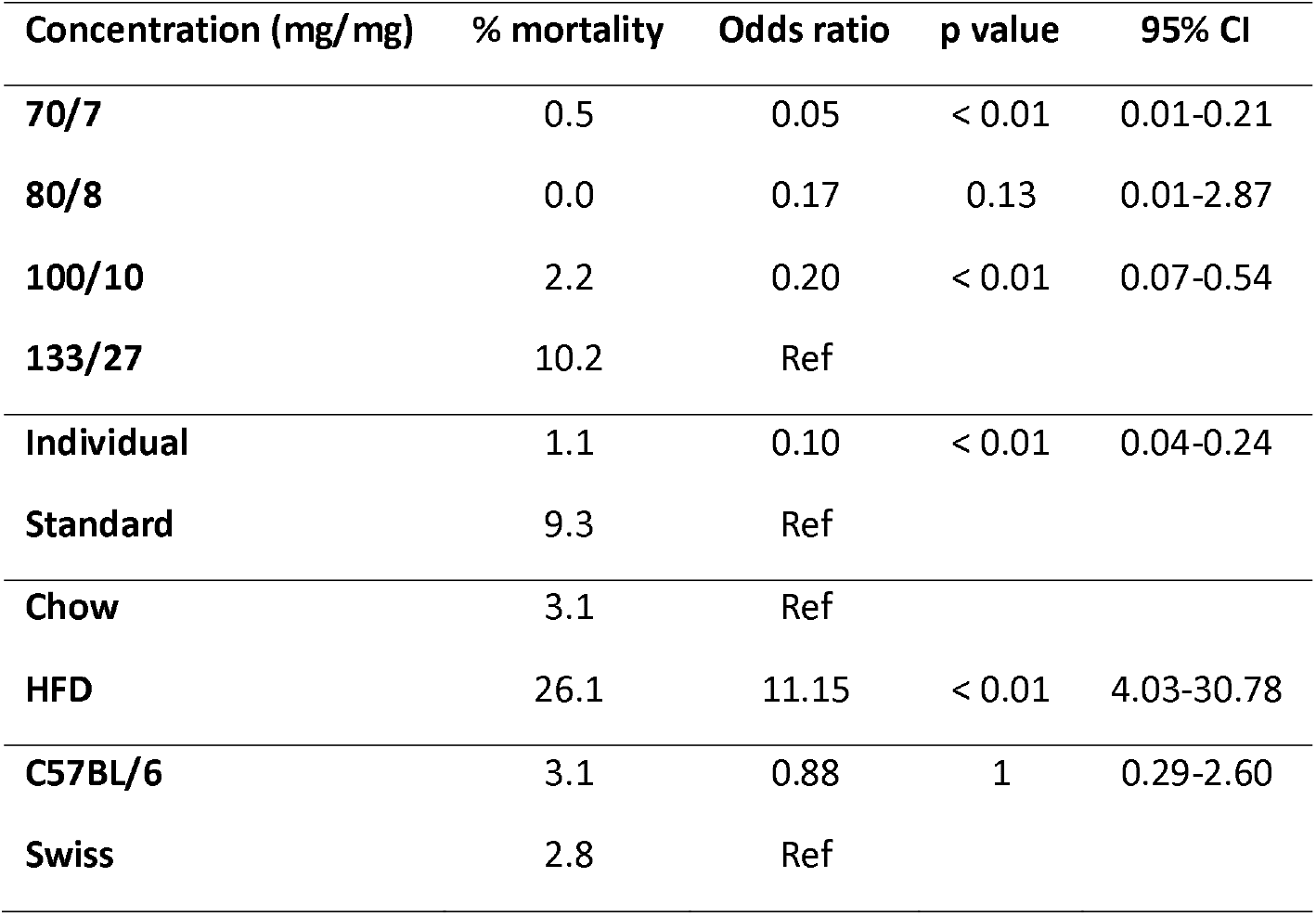
Drug prescription incorporated by Labinsane formulas. The 70/7, 80/8, 100/10 (Labinsane) and 133/27 (standard dose) refers to mg ketamine/mg xylazine for kilograms. Individual dose (weight-adjusted using Labinsane) compared to standard anesthetic dose. Animals fed a Chow diet compared to animals fed a high-fat diet. C57BL/6J mice compared to Swiss mice strain. Fisher’s exact test. The two-tailed p value. CI: Confidence Interval.

## Discussion

Currently, experimental animals are widely used in biological and medical research. However, the scientific community has raised several bioethical concerns, such as the number of animals required to achieve reproducible and statistically relevant results. These concerns involve aspects related to pain, discomfort, and unwanted animal loss during procedures. The Principles of Humane Experimental Technique published in 1959 by Russell and Burch proposed that all efforts should be made to minimize the use and suffering of experimental animals in biological and health research (3R). Today, after 60 years, we are still struggling to achieve the high standards idealized by Russell and Burch.

Despite the recommended anesthetic doses being well-known worldwide, the final anesthetic doses could be different than those calculated. Several animals for experiments, preparation of master anesthetic solutions, small drug volumes and volumetric limitations of syringe systems could interfere in the proper application of recommended dose rates. We asked if we could improve the reported average mortality using ketamine and xylazine. First, we tested a Microsoft Excel spreadsheet containing the proper prescription formula. However, after implementation, adherence to the new weight-adjusted dose brought difficulties for new operators. For instance, aseptic concerns about personal computers in laboratories performing animal procedures, low volumes of anesthetic to remove from the flask, highly concentrated residual anesthetic volumes in the syringe and limitations of syringe systems without the capacity to fractionate small volumes of anesthetic. To solve some of these complexities, we tried a different approach and developed a mobile app. Using the Labinsane mobile app on personal mobile phones, researchers and operators were able to use personal devices in the workplace. Mobile phones were easily disinfected and sped up master anesthetic cocktail calculations, as well as the individual weight-adjusted anesthetic dose calculation.

Several advantages of volatile anesthetics over injectables have been shown. However, anesthetic procedures using airways requires specific equipment, which can interfere with the experiment result and increases both cost and workspace usage. If volatile anesthetics cannot be applied, injectable anesthesia is indicated ^13^. Intraperitoneal anesthetic protocols based on the injection of ketamine-xylazine solution are widely used in experiments with rodents, due to the low cost, minimum training and no equipment being required ^2^. Nevertheless, there is a wide variation in the recommended dose, which is possibly due to differences between mouse strains, type and duration of the procedure, health conditions, age and research goals ^2,6,8,14^. Previous reports compared the efficacy of the intraperitoneal and subcutaneous administration of ketamine (100 mg/ml) and xylazine (20mg/ml) solution for inducing surgical anesthesia. Among C57BL/6, BALB and ICR mice, no deaths occurred in the subcutaneous administration groups, while16.7% (10 of 60 mice) of mice injected intraperitoneally died ^8^. Also, previous reports have shown that the ketamine, xylazine and diazepam cocktail is effective to induce anesthesia for surgical procedures ^15-19^. To improve user experience, we developed a mobile app for use during anesthetic procedures.

The principal causes of unsafe medications in humans are related to failures in drug preparation and lack of treatment standardization ^20^. Experimental animal safety should also ensure the proper anesthetic dose calculation and administration. In this way, the Labinsane mobile app seeks to ensure the safety of C57BL/6 and Swiss mice, pursuing proper individual anesthetic dose calculation and administration.

We decided to use “4” and “3” as the dilution factors in the Labinsane formulas with the aim to increase the total volume to administer. Increasing the dilution factor, we decreased the anesthetic agent concentrations in every microliter of the final cocktail. In this way, if any operator mistake occurs, the amount of anesthetic agents is low (in μL), protecting mice from overdoses. Also, these dilution factors helped operators with anesthetic volumes that were easier to fit in common the insulin syringe (1 ml). Altogether, this protects mice from human mistakes. This approach suggests that the Labinsane mobile app improves mice safety related to anesthetic administration.

The MIT App Inventor platform (appinventor.mit.edu) allows researchers to easily create new mobile apps. Indeed, App Inventor apps have been shown to improve data analysis and help in making well-informed decisions ^10^. Also, the use of the App Inventor has contributed to children’s learning ^21^, home automation ^22^ and self-care actions ^23^. Our study established that Labinsane, a MIT based mobile app, helps researcher to markedly reduce anesthesia-related mortality. We encourage researchers to validate new experimental animal strains and species based on modular collaboration with Labinsane to decrease anesthetic-related death.

Also, we evaluated if two different diet, Chow and HFD, could influence the anesthesia-related mortality rate. In this way, we identified that animals fed a HFD showed the highest anesthesia-related mortality (26.1%). We identified that animals fed a HFD are 11.1 times more likely to suffer intraoperative and anesthesia-related deaths compared to animals fed with Chow (OR 11.1, 95% CI 4.03-30.78, p < 0.01). Consistent with previous reports ^24,25^, our results verified that obese animals had a higher mortality rate compared to normal weight animals. Indeed, obese mice were shown to have a 100% mortality after surgery, even with no evident medical complications ^25^. In the same way, our data showed 26% mortality in obese HFD animals after surgery. This could be explained because obese mice have larger deposits of adipose tissue ^26^, alterations in binding, and distribution and excretion properties of anesthetics, finally impacting on the pharmacokinetics and pharmacodynamics of anesthetic agents. However, future studies will be necessary to determine new strategies to avoid obesity-related mortality in anesthetized mice.

## Acknowledgements

The authors are grateful to Marcio Alves da Cruz, Joseane Morari, Erika Anne Roman and Gerson Ferraz for technical assistance.

This study was financed in part by the Coordenação de Aperfeiçoamento de Pessoal de Nível Superior – Brasil (CAPES) – Finance Code 88882.434714/2019-01.

## Author contributions

All the authors reviewed and approved the manuscript.

