## Supplementary material for "A smartphone app for individual anesthetic calculation decreased anesthesia-related mortality in mice": Table 2

|  | **Excel formula** | **Labinsane** |
| --- | --- | --- |
| **Number animals** | 449 | 449 |
| **Minimum (μL)** | 114.6 | 114.6 |
| **Median (μL)** | 130.7 | 130.7 |
| **Maximum (μL)** | 249.8 | 250 |
| **Mean (μL)** | 146.1 | 145.9 |
| **Std. Deviation (μL)** | 31.9 | 31.6 |
| **Std. Error of Mean (μL)** | 1.5 | 1.5 |
| **Lower 95% CI (μL)** | 143 | 143 |
| **Upper 95% CI (μL)** | 149 | 148.8 |

Table 2. Labinsane validation. Median, mean, standard deviation, standard error and confidence interval comparing the Microsoft Excel spreadsheet and Labinsane mobile app. Unpaired t test, p value: 0.9432.
