## Supplementary material for "A smartphone app for individual anesthetic calculation decreased anesthesia-related mortality in mice": table 3

| **Concentration (mg/mg)** | **% mortality** | **Odds ratio** | **p value** | **95% CI** |
| --- | --- | --- | --- | --- |
| **70/7** | 0.5 | 0.05 | < 0.01 | 0.01-0.21 |
| **80/8** | 0.0 | 0.17 | 0.13 | 0.01-2.87 |
| **100/10** | 2.2 | 0.20 | < 0.01 | 0.07-0.54 |
| **133/27** | 10.2 | Ref |  |  |
| **Individual** | 1.1 | 0.10 | < 0.01 | 0.04-0.24 |
| **Standard** | 9.3 | Ref |  |  |
| **Chow** | 3.1 | Ref |  |  |
| **HFD** | 26.1 | 11.15 | < 0.01 | 4.03-30.78 |
| **C57BL/6** | 3.1 | 0.88 | 1 | 0.29-2.60 |
| **Swiss** | 2.8 | Ref |  |  |

Table 3. Drug prescription incorporated by Labinsane formulas. The 70/7, 80/8, 100/10 (Labinsane) and 133/27 (standard dose) refers to mg ketamine/mg xylazine for kilograms. Individual dose (weight-adjusted using Labinsane) compared to standard anesthetic dose. Animals fed a Chow diet compared to animals fed a high-fat diet. C57BL/6J mice compared to Swiss mice strain. Fisher's exact test. The two-tailed p value. CI: Confidence Interval.
