## Supplementary material for "A smartphone app for individual anesthetic calculation decreased anesthesia-related mortality in mice": Table 1

|  | **Survival** | **Non-survival** | **Total** |
| --- | --- | --- | --- |
| **Individual dose** | 610 | 7* | **617** |
| **Standard dose** | 149 | 17 | **166** |
| **Total** | **759** | **24** | **783** |

Table 1. Contingency table. Survival and non-survival numbers crossed by Individual dose (weight-adjusted dose) and standard dose in Chow diet animals. Fisher's exact test. * p < 0.05
